# Glycocalyx-induced formation of membrane tubes

**DOI:** 10.1101/2024.11.27.625577

**Authors:** Ke Xiao, Padmini Rangamani

**Author notes:** To whom correspondence must be addressed.

## Abstract

Tubular membrane structures are ubiquitous in cells and in the membranes of intracellular organelles such as the Golgi complex and the endoplasmic reticulum. Tubulation plays essential roles in numerous biological processes, including filopodia growth, trafficking, ion transport, and cellular motility. Understanding the fundamental mechanism of the formation of membrane tubes is thus an important problem in the fields of biology and biophysics. Though extensive studies have shown that tubes can be formed due to localized forces acting on the membrane or by the curvature induced by membrane-bound proteins, little is known about how membrane tubes are induced by glycocalyx, a sugar-rich layer at the cell surface. In this work, we develop a biophysical model that combines polymer physics theory and the Canham-Helfrich membrane theory to investigate how the glycocalyx generates cylindrical tubular protrusions on the cell membrane. Our results show that the glycocalyx alone can induce the formation of tubular membrane structures. This tube formation involves a first-order shape transition without any externally applied force or other curvature-inducing mechanisms. We also find that critical values of glycocalyx grafting density and glycopolymer length are needed to induce the formation of tubular structures. The presence of vertical actin force, line tension, and spontaneous curvature reduces the critical grafting density and length of polymer that triggers the formation of membrane tube, which suggests that the glycocalyx makes tube formation energetically more favorable when combined with an actin force, line tension, and spontaneous curvature.

**Significance Statement:** In many cells, the existence of glycocalyx, a thick layer of polymer meshwork comprising proteins and complex sugar chains coating the outside of the cell membrane, regulates the formation of membrane tubes. Here, we propose a theoretical model that combines polymer physics theory and the Canham-Helfrich membrane theory to study the formation of cylindrical tubular protrusions induced by the glycocalyx. Our findings indicate that glycocalyx plays an important role in the formation of membrane tubes. We find that there exists critical grafting density and length of polymer that triggers the formation of membrane tubes, and the glycocalyx-induced tube formation is facilitated when combined with actin forces, line tension, and spontaneous curvature. Our theoretical model has implications for understanding how biological membranes may form tubular structures.

## Introduction

The generation of elongated tubular membrane geometries is ubiquitous in both plasma membrane and organelle membranes, including the endoplasmic reticulum (*1, 2*), the Golgi apparatus (*3, 4*), and the inner mitochondrial membrane (*5*–*7*). Such elongated tubular structures play essential roles in numerous biological processes ranging from membrane trafficking to ion transport and cellular motility (*8*–*11*). Thus, understanding the fundamental mechanism of the formation of tubular membrane geometries with high curvature is an important problem in the fields of biology and biophysics.

In *in vitro* experiments, cylindrical tubes can be created using various experimental techniques including hydrodynamic flow (*12*), micropipette aspiration (*13*), and a mix of micromanipulation and optical (*14*) or magnetic (*15*) tweezers. These approaches can be attributed to a pulling force exerted on a localized point on the membrane. *In vivo*, additional mechanisms to induce the generation of elongated protrusions such as forces acting on the membrane by cytoskeletal assembly, filament bundles, and motor proteins (*10, 16*–*21*) or the interaction of cellular membrane with intrinsically curved proteins and oligomers (*22*) or curvature-inducing proteins (i.e., BAR(Bin/Amphiphysin/Rvs)-domain proteins) without any apparent localized force mechanisms (*23*–*26*) also generate tubular structures. Recently, a wide range of studies have observed that many protrusive membrane structures such as epithelial microvilli (*27*–*32*), cilia (*33*), and filopodia (*34, 35*) can be generated by mucins, a class of large, heavily glycosylated proteins that partially make up the glycocalyx. The glycocalyx is a thick layer of heavily glycosylated transmembrane macromolecules concentrated on most cell surfaces in a complex brush structure (*36*–*39*). Although many experimental studies indicate that the glycocalyx plays an essential role in the formation of membrane tubes, theoretical models that can explain the force generation mechanisms of the glycocalyx on the cell surface remain underdeveloped.

Experimentally, to explore the mechanisms of membrane shape regulation by the glycocalyx, Shurer et al. (*39*) reported that bulky brush-like glycocalyx polymers are sufficient to induce a variety of curved membrane features, including spherical-shaped membranes (referred to as blebs), tubes, and unduloids, in a density-dependent manner. From a theoretical point of view, the elastic continuum models based on the Canham-Helfrich theory (*40, 41*) are frequently used to investigate the formation of membrane tubes. Meanwhile, through utilizing mean-field theory, analytical methods (*42*–*45*), scaling theory (*46*), phase field model (*47, 48*), and Monte Carlo simulations (*49, 50*), theoretical studies have shown that anchored polymers on membrane give rise to the changes of membrane shape, membrane elastic properties, and its spontaneous curvature. Recently, we developed a general energetic framework that couples the mechanics of the glycocalyx with the mechanics of the lipid bilayer to investigate how the glycocalyx can generate spherical vesicles and to explore whether the glycocalyx itself can sense curvature (*51*). However, the theoretical frameworks that can describe the possible underlying mechanisms of glycocalyx-induced formation of membrane tubes remain lacking, restricting our ability to predict the effects of glycocalyx properties (grafting density and length of glycocalyx) on tubular membrane structures.

In this work, we developed a theoretical model for glycocalyx-induced membrane deformation based on polymer physics theory and the Canham-Helfrich membrane theory. Our model includes the effects of glycocalyx properties, actin force, and membrane properties on the formation of tubular membrane structures. We systematically investigated the influence of different factors on the formation of membrane tubes. Our model predicts that the properties of the glycocalyx are a key determinant in inducing tubular membrane structures. Membrane tubes can be induced by glycocalyx when the glycocalyx grafting density and glycopolymer length exceed a threshold value. Denser and longer glycopolymers on the membrane surface promote the formation of longer and thinner tubular structures. The corresponding critical values of the glycocalyx properties are reduced in the presence of actin force, line tension, and spontaneous curvature. This work provides a unifying framework for membrane shape remodeling by integrating multiple related factors and provides fundamental insight into the regulation of tubular structures.

## Model and Methods

### Description of the model

Here, we develop a biophysical model for the glycocalyx-membrane composite, where a layer of glycocalyx is coated on a membrane surface (Fig. 1 (a, b)). A recent study has shown that the entropic forces generated by the glycocalyx on the outer surface of the cell membrane can induce finger-like extensions (*39*). To investigate the effect of glycocalyx on the generation of tubular membrane morphology (Fig. 1 (c)), we assume a membrane domain anchored by glycocalyx biopolymers with fixed surface area 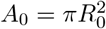 is deformed into a family of cylindrical tubes with length *L*_*t*_ connected with a hemispherical cap with radius *R*_*t*_, where *R*_0_ is the in-plane radius of the flat membrane, as shown in Fig. 1(d, e). In the glycocalyx-membrane system, we assume that the layer of glycopolymers is uncharged and in a brush-like structure, where the polymer brushes are extended equally in height, *L*_brush_. We define a shape parameter, *χ* = *L*_*t*_*/R*_*t*_, which characterizes the tube shape. Then, the radius of the tube can be expressed as 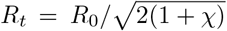 under the constraint of fixed polymer grafting area 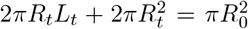. When *χ* = 0, the formation of a tubular shape is unfavorable. The membrane domain takes the shape of a family of tubes with a hemispherical cap connected when *χ* ≠ 0. The energy of the polymer layer depends on the polymer properties including the grafting density, length of the polymer, and the interactions between the individual polymer chains, and also on the curvature of the underlying substrate (*52*). Our approach is to find the shape that corresponds to the minimal energy state for the combined energy of the membrane and the polymer for cylindrical deformations.

**Figure 1:**
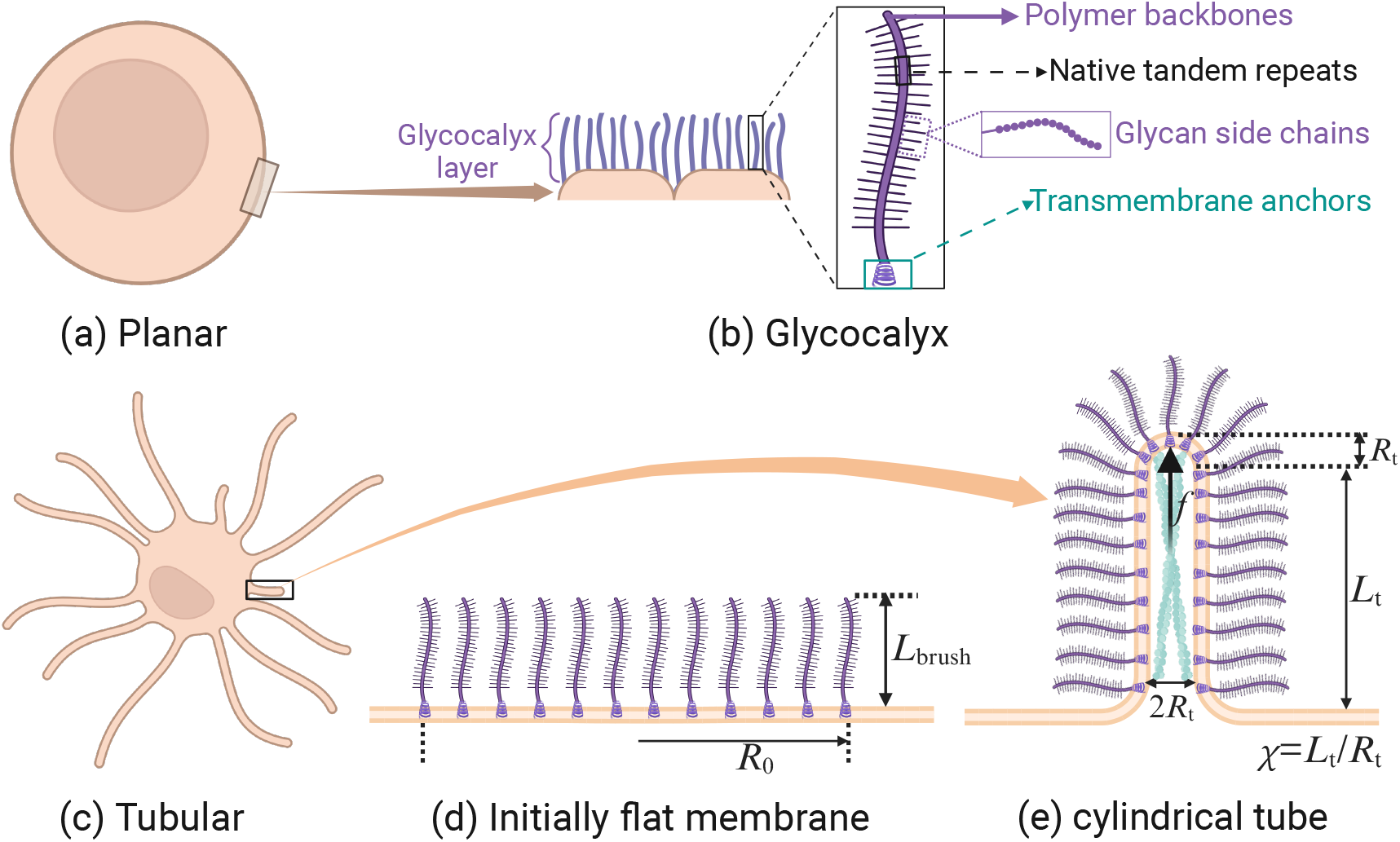
Schematic of glycocalyx-membrane system. Schematic showing (a) a mostly flat cell membrane surface with (b) a sugar-rich layer of glycocalyx coating on its outer surface. An enlarged view of the polymer structure of glycocalyx constituents such as mucins. (c) Sketch of tubular membrane morphology generated by the glycocalyx. (d) An initially flat bilayer grafted with a patch of glycocalyx of radius *R*_0_ and brush height *L*_brush_. (e) An enlarged geometrical sketch of the system: a cylindrical tube of radius *R*_*t*_ and length *L*_*t*_, with a hemispherical cap of radius *R*_*t*_. The light blue rod inside the tube represents the cartoon schematic of a filamentous actin (F-actin) core.

### Energetics of the system

The total free energy of the glycocalyx-membrane system (*F*_tot_) is a sum of four terms: the energy contribution associated with the glycocalyx (*F*_glycocalyx_), the elastic energy of the membrane (*F*_membrane_), the line tension energy (*F*_line tension_), and the work done by the vertical actin force (*F*_force_)

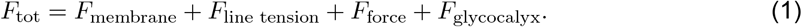

The lipid bilayer is modeled as a thin elastic shell, which resists out-of-plane bending and we assume that the Helfrich energy is sufficient to capture membrane bending. The energy cost of bending a membrane is given by the well-known Canham-Helfrich Hamiltonian (*40*) which includes the bending energy of the membrane and the surface tension energy. Thus, the elastic energy of the membrane appearing in Eq. (1) can be written as

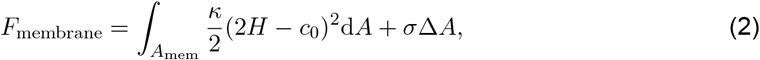

where *κ* is the membrane bending rigidity, *H* = (*c*_1_ + *c*_2_)*/*2 is the mean curvature in which *c*_1_ and *c*_2_ are the two principal curvatures, *c*_0_ is the spontaneous curvature, *σ* is the membrane tension, and Δ*A* is the change in the in-plane area due to the membrane shape deformation. Here, *c*_0_ is the spontaneous curvature which is restricted to asymmetries in the bilayer.

To take into account the multiple phases on the membrane, for example, the glycocalyx rich domain and the surrounding bare membrane, we also considered the effect of the line tension at the glycopolymers anchored domain boundary or interface *∂l*. Therefore, the second term on the right-hand side of Eq. (1) is the interfacial energy or the energy associated with the line tension and is given by

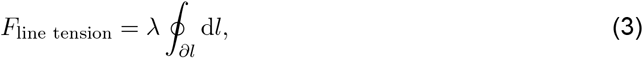

where *λ* denotes the strength of line tension along the domain boundary, and the integral is over the periphery line d*l* of the domain. The integral is over the periphery line d*l* of the membrane patch on which the glycocalyx is anchored. We assume that the value of line tension is a constant. Intuitively, this line tension describes the energetic cost of maintaining a high density and a low density region.

The third term on the right-hand side of Eq. (1) describes the work done by the vertical actin force. The polymerization of actin filaments can generate pushing force for protrusion in living cells (*18, 53, 54*). Notably, Ref. (*39*) showed that the polymerization and depolymerization of cytoskeletal filamentous actin (F-actin) core (see Fig. 1(e)) also plays a key role in mediating membrane shape. Because the precise architecture formed by the F-actin core inside the cell and the resulting forces are not yet well established, so we assume that the F-actin cores form bundles (rods) that apply vertical forces on the membrane. Note that we assume the vertical force is a point force (*21, 55*–*57*) in the *z* direction in our model. Based on such an assumption, we model the force generated by the F-actin core, *f*, as

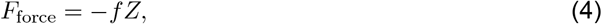

where *Z* is the height of the deformed membrane patch.

We restrict the membrane geometry to a cylindrical tube connected with a spherical cap. The two principal curvatures for the cylindrical tube are given by *c*_1_ = 0 and *c*_2_ = 1*/R*_*t*_, and for the spherical cap are given by *c*_1_ = *c*_2_ = 1*/R*_*t*_. As a result, the excess area is calculated as 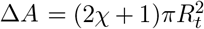, the length of the domain boundary can be written as ∮_*∂l*_ d*l* = 2*πR*, and the height of the deformed membrane patch is given by *Z* = *R*_*t*_ + *L*_*t*_. Therefore, the sum of the elastic energy of the membrane, the line tension energy, and the work done by the vertical actin force becomes

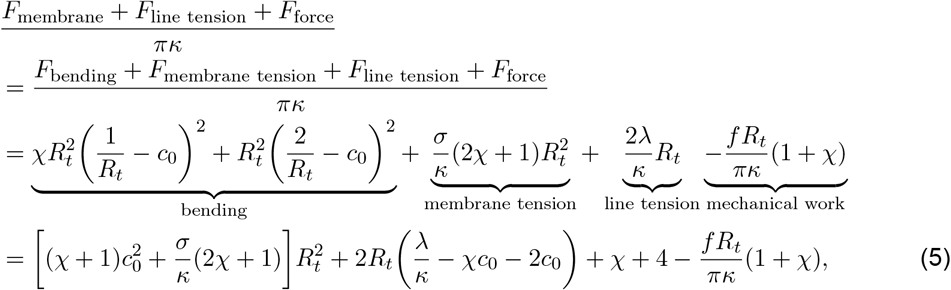

where dividing by *πκ* normalizes the energy.

The final term in Eq. (1) is the energy contribution associated with the glycocalyx, which includes the elastic stretching of the polymer chains and the interactions between monomers in the polymer brush. Based on Ref. (*52*), the sum of the energy density of these two terms for a single polymer is given by

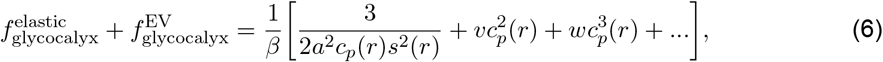

where *β* = 1*/*(*k*_*B*_*T*) with *k*_*B*_*T* being the unit thermal energy, *a* is the monomer size, *c*_*p*_(*r*) is the local monomer density profile along the thickness, *r* is the radial distance defined from the center of the spherical or cylindrical surface, *s*(*r*) is the area per chain at distance from the polymer grafting surface, and *υ* and *w* are the second and third virial coefficient, respectively. Integrating this energy density from *R*_*t*_ to *R*_*t*_ + *L*_brush_ and then multiplying the total number of grafted polymers for the cylindrical tube and the hemispherical cap, respectively, leads to the total energy contribution from glycocalyx polymers

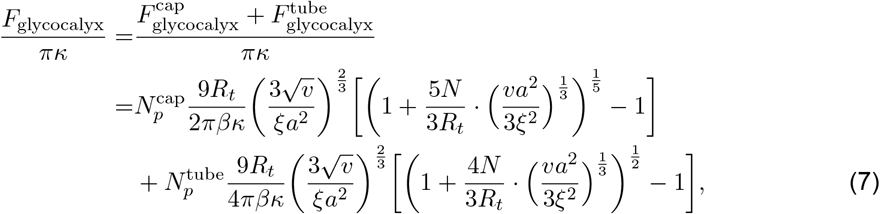

where 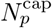 and 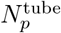 are the number of polymer chains that grafted on the spherical cap and the cylindrical tube, *ξ* is the grafting distance which is related to the grafting density *ρ* via *ρ* = 1*/ξ*^2^, and *N* is the number of monomers in a polymer chain. Here, the relation *χ* = *L*_*t*_*/R*_*t*_ is used, and the detailed derivation of this individual energy component (Eq. (7)) is provided in the Supplemental Material. Note that only the pairwise monomer-monomer interactions with second virial coefficient *υ* is considered, and that the ternary interactions with third virial coefficient *w* is neglected.

Finally, combining Eqs. (5) and (7), and using 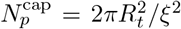 and 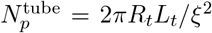, the total free energy of the glycocalyx-membrane system with a cylindrical tube of radius *R*_*t*_ and length *L*_*t*_, and with a hemispherical cap of radius *R*_*t*_ can be obtained as

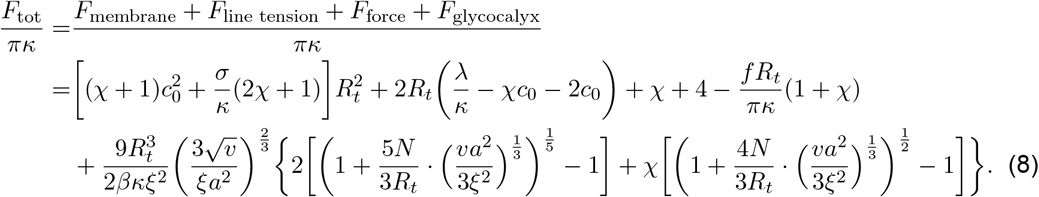

Hereafter, in our model, we use the number of monomers, *N*, to capture the length of the polymer due to that the thickness of the polymer brush is related to the total number of monomers *N* (see Eqs. (S9) and (S12) in the Supplemental Material). We assume that our system is at equilibrium, implying that the system selects the membrane shape that minimizes the total free energy of Eq. (8).

### Numerical implementation

Here, we are focused on the equilibrium state of the system, so our objective is to determine the global minimum energy state by numerically calculating the corresponding total free energy of the system. To numerically calculate the total free energy, the excluded volume parameter is set as *υ* = *a*^3^ as suggested by (*52*) for neutral brushes. We have combined these contributions from the glycocalyx polymers and the cell membrane to construct the total free energy of the membrane patch with fixed area *A*_0_. Minimization of the total energy as a function of shape parameter *χ* yields the optimal *χ*_min_ that determines the morphology of the membrane patch. Biologically relevant values for the parameters that have been used in the mathematical model are summarized in Tables 1. The code is available on [https://github.com/RangamaniLabUCSD/Glycocalyx_membrane-tubes].

**Table 1:**
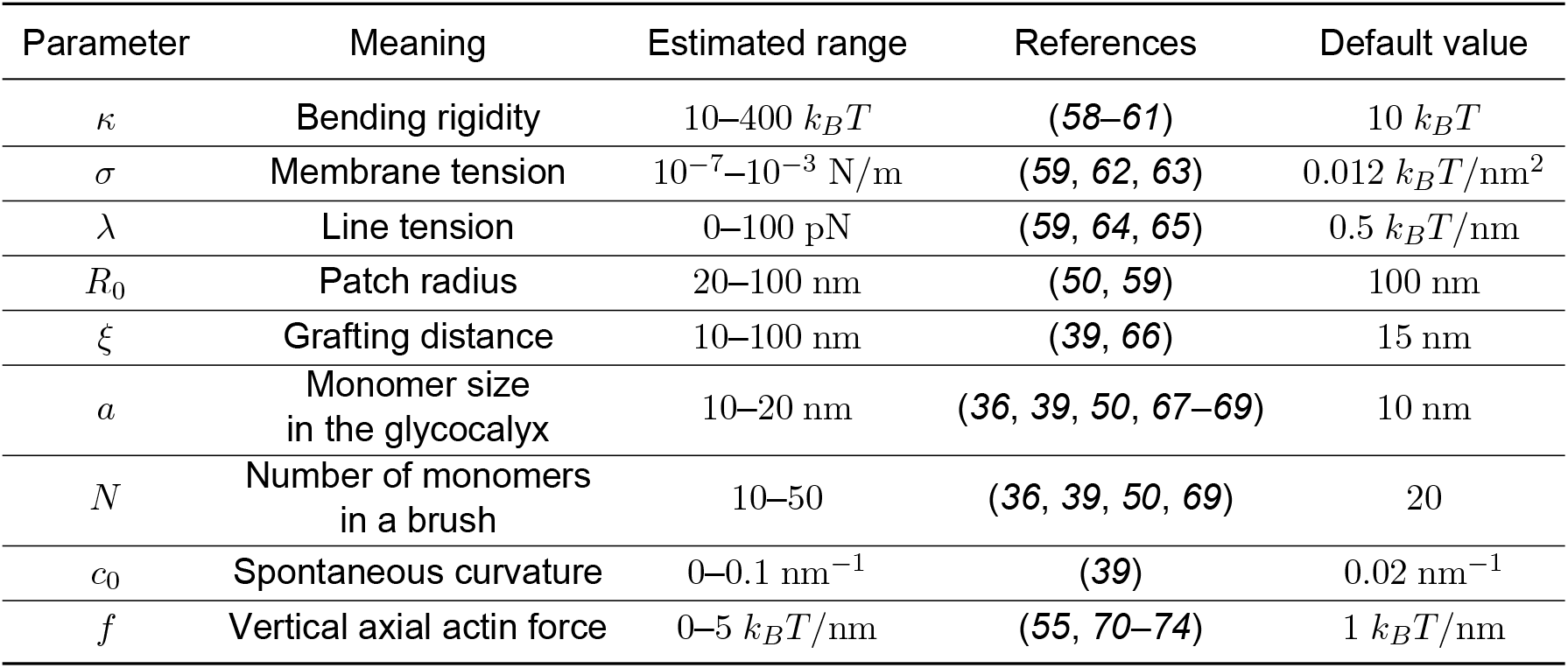
Values of the different parameters used in the model.

## Results

In order to unveil the effects of glycocalyx properties, membrane properties, and actin force on the formation of membrane tubular, we numerically calculate the total energy for different shape parameters *χ* by varying each of such parameters one by one. The global energy minimum corresponds to the optimal *χ*_min_ that gives the equilibrium geometry of the glycocalyx-covered membrane patch. We used default parameter values listed in Table 1, unless otherwise indicated.

### Effect of glycocalyx on membrane tubular formation

To investigate the effects of the glycocalyx properties including grafting density and length, we first perform the analysis of total energy for different values of grafting density *ρ* and length *N*, as shown in Fig. 2. In Fig. 2(a), we plot the total energy as a function of the shape parameter *χ* for different grafting densities. Here, the corresponding contributions from different energy components are plotted in Fig. S2. Note that the total energy is a nonlinear function of the shape parameter. The energy curve shows a monotonic behavior when the glycopolymer grafting density is smaller than a critical value (see the purple curve *ρ* = 0.0047 nm^*−*2^), where there is only one local minimum (global minimum) at *χ* = 0 (solid circle in the purple curve), corresponding to zero tube length. This indicates that when the glycocalyx is sparsely distributed on the membrane, tube formation is not energetically favorable. As grafting density increases, the energy profile corresponds to two local minima: one stable non-tubular state (*χ* = 0), and another metastable tubular state (*χ*≠ 0). This suggests that the initial spherical cap shape is still more favorable as compared to the tubular structure (see the green and blue curves). If the grafting density is larger than a critical value (*ρ* = 0.0077 nm^*−*2^), the total energy of the non-tubular state becomes equal to that of the tubular state (see the red curve), indicating that the two states coexist. Further increase of grafting density leads to the transition from a non-tubular state to a tubular state (see the black curve), which implies that the initial spherical cap becomes unstable against the formation of a tube. Note that the initial spherical cap, in order to grow up to its preferred length (solid point on the black curve), needs to overcome an energy barrier Δ*F*. This observation is consistent with the conclusion of previous studies (*48*). Campelo et al. (*48*) argue that polymer-induced tubulation is caused by the nonhomogeneous concentration of amphiphilic molecules anchored on the membrane. However, the energy barrier in our model comes from the nonlinear dependence of the energy contribution associated with the glycocalyx on the shape parameter.

**Figure 2:**
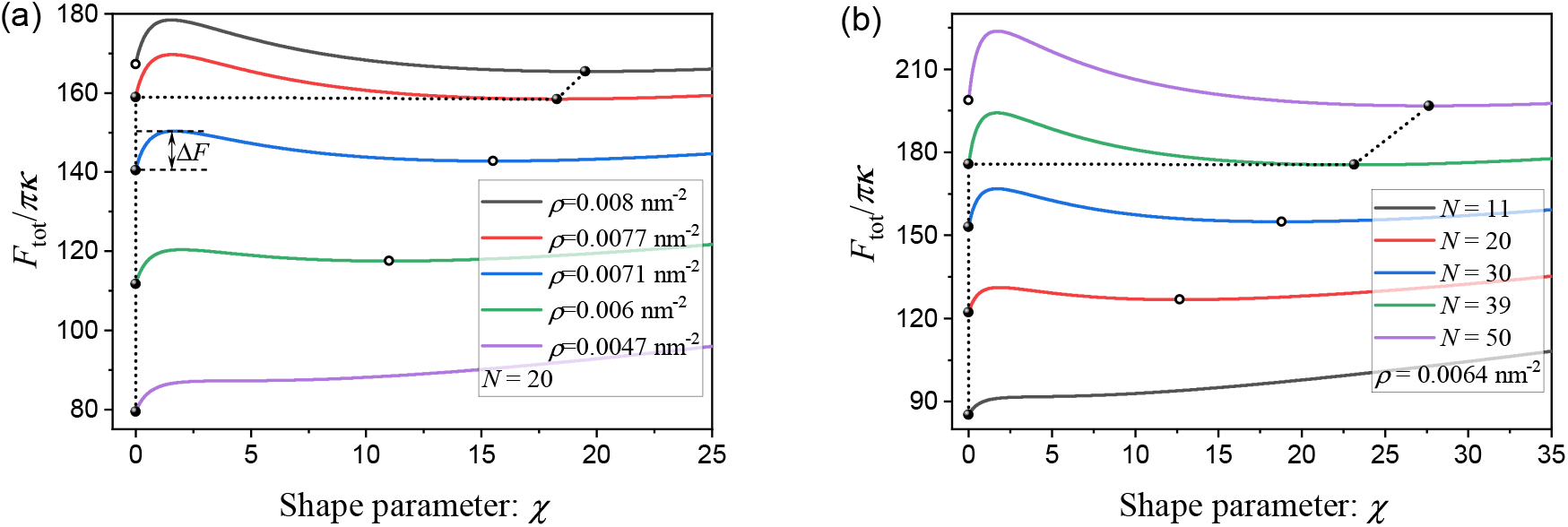
Total energy profile as a function of shape parameter *χ* for different values of (a) glycopolymer density *ρ* and (b) glycopolymer length *N*, where the bending modulus, membrane tension, line tension, spontaneous curvature, and vertical force are fixed at *κ* = 10 *k*_*B*_ *T, σ* = 0.012 *k*_*B*_ *T/*nm^2^, *λ* = 0 *k*_*B*_ *T/*nm, *c*_0_ = 0 nm^*−*1^, and *f* = 0 *k*_*B*_ *T/*nm, respectively.

In addition to grafting density, length of the polymers is another important feature of the glycocalyx layer on the cell membrane surface. To probe the influence of glycopolymer length on the formation of membrane tube, the analysis of total energies for various numbers of monomer *N* is presented in Fig. 2(b). Upon increasing *N*, the energy profiles in Fig. 2(b) show a similar trend as presented in Fig. 2(a), indicating that longer glycopolymer length can trigger a state transition from non-tubular to tubular (i.e., the purple curve *N* = 50). Thus, the glycopolymer length also has a big impact on the formation of membrane tubes. The finding of the tube shape induced by the glycocalyx is in line with those observed experimentally by Shurer et al. (*39*).

### Discontinuous transition from non-tubular state to tubular state

We next focused on the stable state corresponding to the shape parameter *χ*_min_. Figure. 3 shows the dependence of *χ*_min_ on grafting density and glycopolymer length. In the absence of line tension *λ*, spontaneous curvature *c*_0_, and actin force *f*, the black dotted curves in Figs. 3(a) and 3(b) illustrate that the optimal shape parameter shows a sharp jump at the critical values, *ρ*_*c*_ or *N*_*c*_, a characteristic of discontinuous transition. Such a transition indicates that the glycocalyx alone is able to induce the membrane to form a stable tubular morphology. Furthermore, to investigate the influences of line tension *λ*, spontaneous curvature *c*_0_, and actin force *f* on the dependence behaviors of *χ*_min_ on glycopolymer density *ρ* and length *N*, the orange and blue curves in Figs. 3(a) and 3(b) show that the discontinuous transition is still maintained but with a shift of the threshold *ρ*_*c*_ or *N*_*c*_ which triggers the discontinuous transition from non-tubular state to tubular state. Moreover, we found that the presence of line tension *λ*, spontaneous curvature *c*_0_, and actin force *f* results in the decreases of the critical value *ρ*_*c*_ or *N*_*c*_, as shown in Figs. 3(c) and 3(d). Also, we can generate longer and thinner membrane tubes (larger *χ*_min_ = *L*_*t*_*/R*_*t*_) in the presence of line tension *λ*, spontaneous curvature *c*_0_, and actin force *f*. Consequently, we can infer that the presence of line tension, spontaneous curvature, and actin force plays a role in relaxing the critical conditions for the formation of membrane tubes.

**Figure 3:**
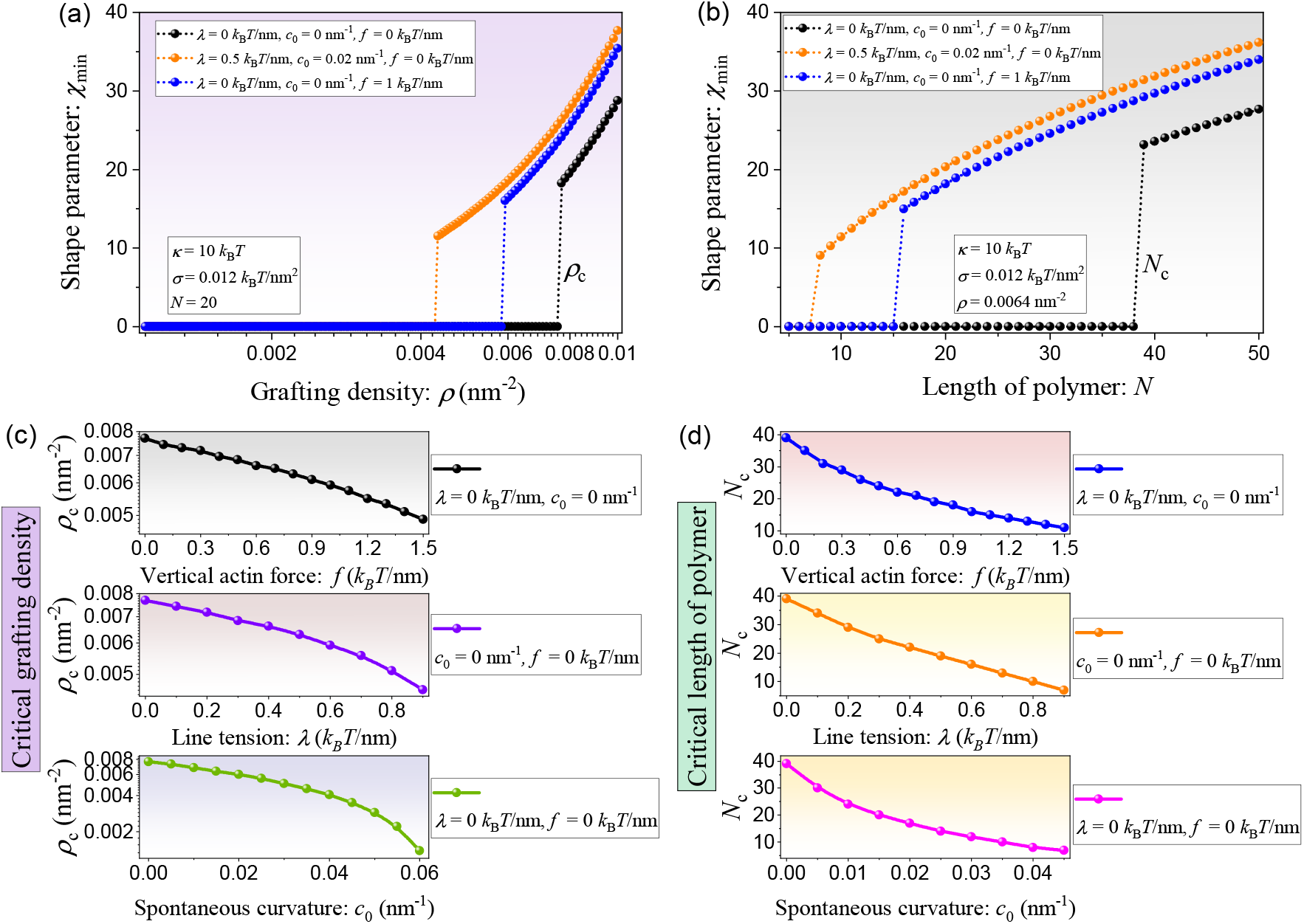
The dependence of minimum shape parameter *χ*_min_ on glycopolymer density *ρ* with *N* = 20 and (b) glycopolymer length *N* with *ρ* = 0.0064 nm^*−*2^, where the bending modulus and membrane tension are fixed at *κ* = 10 *k*_*B*_*T* and *σ* = 0.012 *k*_*B*_*T/*nm^2^, respectively. The dependence of (c) critical grafting density, *ρ*_*c*_, and (d) critical length of polymer, *N*_*c*_, on actin force (top row), line tension (middle row), and spontaneous curvature (bottom row).

### Phase diagrams for glycocalyx-induced tubular formation

In order to systematically study the interrelated effects of glycopolymer density and length on the membrane shape, we summarize the observed stable states in phase diagrams presented in Fig. 4 for four different cases: (i) in the absence of line tension *λ*, spontaneous curvature *c*_0_, and actin force *f* (see Fig. 4(a)); (ii) in the presence of actin force *f* and in the absence of line tension *λ* and spontaneous curvature *c*_0_ (see Fig. 4(b)); (iii) in the absence of actin force *f* and in the presence of line tension *λ* and spontaneous curvature *c*_0_ (see Fig. 4(c)); and (iv) in the presence of line tension *λ*, spontaneous curvature *c*_0_, and actin force *f* (see Fig. 4(d)). Figure. 4 shows that two regimes corresponding to non-tubular and tubular states are identified, in which the color indicates the value of the optimal shape parameter *χ*_min_. The discontinuous transition from a non-tubular state to a tubular state can be triggered by tuning the grafting density and length of the glycocalyx. Fig. 4(a) revealed that, even without the line tension, spontaneous curvature, and actin force, the glycocalyx alone can regulate the formation of tubular structures under the condition of crossing the threshold of the grafting density and polymer length. When comparing Fig. 4(a) with Fig. 4(b), we observe that the inclusion of vertical actin force leads to a broader tubular regime and shifts the boundary curve to the left (see the orange curve). This validates the prediction presented in the top row in Figs. 3(c) and 3(d), which is that lower *ρ*_*c*_ or *N*_*c*_ is sufficient to induce the formation of membrane tube when other factors are present. Furthermore, the comparison of the tubular region between Fig. 4(a) and Fig. 4(c) shows that the tubular regime becomes wider in the presence of line tension and spontaneous curvature. This confirms the predictions illustrated in the middle and bottom rows of Figs. 3(c) and 3(d), which shows that the threshold for inducing the formation of tubules decreases as the line tension and spontaneous curvature increase. In the end, a direct comparison of the tubular region in Fig. 4(d) to that of Fig. 4(a) (or Fig. 4(b) and 4(c)) implies that the line tension, spontaneous curvature, and actin force facilitate the formation of tubules in conditions that would not be favored otherwise. Figure. 4(d) also shows that the unfavorable conditions (no tubular regime) vanish in the selected parameter space when line tension, spontaneous curvature, and actin force are taken into account. We expect that a regime unfavorable to tube formation will emerge when the polymer grafting density and polymer length are sufficiently low.

**Figure 4:**
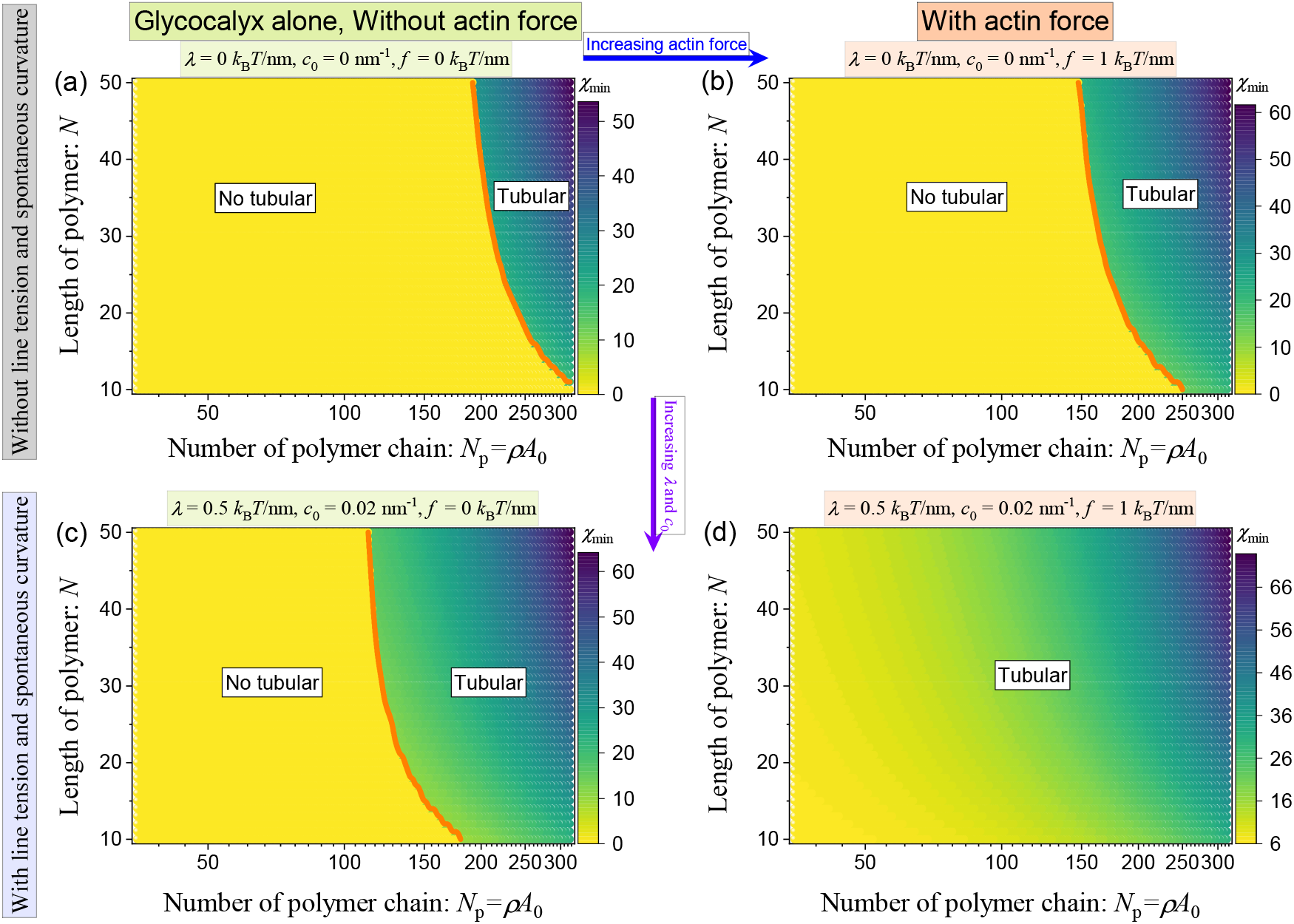
Phase diagrams present the optimal shape parameter *χ*_min_ as a function of number of polymer chain, *N*_*p*_, and length of polymer, *N*, for the situations of (a) without line tension *λ* = 0 *k*_*B*_*T/*nm, spontaneous curvature *c*_0_ = 0 nm^*−*1^, and vertical actin force *f* = 0 *k*_*B*_*T/*nm; (b) with actin force *f* = 1 *k*_*B*_*T/*nm, and without line tension *λ* = 0 *k*_*B*_*T/*nm and spontaneous curvature *c*_0_ = 0 nm^*−*1^; (c) with line tension *λ* = 0.5 *k*_*B*_*T/*nm and spontaneous curvature *c*_0_ = 0.02 nm^*−*1^, and without vertical actin force *f* = 0 *k*_*B*_*T/*nm; (d) with line tension *λ* = 0.5 *k*_*B*_*T/*nm, spontaneous curvature *c*_0_ = 0.02 nm^*−*1^, and vertical actin force *f* = 1 *k*_*B*_*T/*nm.

In general, we find the existence of two stable states in which the system can reside: in the non-tubular state, the initial spherical cap cannot grow into a tubular shape; in the tubular state, a stable tube can be formed after crossing an energy barrier in the presence of glycocalyx. In our theoretical model, the total free energy Eq. (8) includes the contribution of the elastic energy of the membrane, which indicates that membrane shape remodeling is also impacted by membrane properties such as bending rigidity and membrane tension. This prompts us to explore the effects of the membrane properties on tube formation. The corresponding two-dimensional phase diagram on the (*κ*–*σ*) plane to characterize the interrelated effects of membrane bending rigidity and membrane tension on the membrane shape is presented in Figure S3(a). Also, to intuitively observe the influence of line tension, spontaneous curvature, and vertical actin force on tube formation, two more phase diagrams on the (*λ*–*c*_0_) and (*f* –*c*_0_) planes are constructed, as shown in Fig. S3(b) and S3(c). Our results demonstrate that tubule formation is more easily achieved under conditions of lower membrane bending rigidity and tension. Moreover, the presence of line tension, spontaneous curvature, and actin force not only relaxes the critical conditions necessary for the formation of tubular structures but also promotes the generation of longer and thinner tubes.

## Discussion

By combining polymer physics theory and Helfrich membrane theory, here we proposed a theoretical model to elucidate the physical origin behind the glycocalyx-regulated membrane tube formation. Consequently, we argue that glycocalyx plays a significant role in generating membrane tubes. To do so, critical values of grafting density and polymer length need to be reached to induce a tubular membrane by the glycocalyx. The presence of line tension, spontaneous curvature, and actin force can then further promote the formation of membrane tubes. The physics behind the transition between the non-tubular state and tubular state stems from the competition among the glycocalyx energy, the elastic energy (consisting of bending energy and tension energy), and the work done by the vertical actin force. Membrane bending is facilitated by forces due to the glycocalyx and actin, line tension, and spontaneous curvature, whereas membrane elasticity counteracts its deformation.

In summary, based on the developed theoretical framework, we study the membrane tube formation regulated by glycocalyx and find that the presence of glycocalyx is able to trigger a discontinuous non-tubular to tubular transition. Such a transition provides a deeper insight into the membrane tube formation behaviors induced by glycocalyx. We identify that the tube length in the tubular regime can be increased upon regulating many parameters, such as increasing glycocalyx grafting density and length, line tension and spontaneous curvature, and actin force, reducing membrane bending rigidity and tension. Therefore, aside from the polymerization of actin filament bundles against a membrane, the interaction of cellular membranes with proteins that induce curvature, and a pulling force exerted on a membrane, the presence of the glycocalyx, a sugar-rich layer at the cell surface, is another different necessary biophysical condition for the formation of tubular membrane protrusions, particlularly on cellular membranes.

So far, numerous theoretical models have been developed to gain a better understanding of the formation of tubular structures generated by various mechanisms (*18, 21, 24*–*26, 48, 75*). However, our model differs from other models in the following aspects. In our model, tubulation can be generated without a directed force acting on the membrane when it is coated with a layer of glycocalyx, in contrast to previously reported cases (*18, 21, 75*). In addition to the point-like force applied to the membrane, several studies have shown that tubular shapes can also be induced by anisotropic curvature-inducing proteins coating on the membrane (*24*–*26*). In comparison to that protein scaffolding mechanism, here we show a novel mechanism of tube extraction that bulky brush-like glycocalyx polymers are sufficient to induce the formation of a cylindrical tube in the absence of any curvature-inducing proteins. Of note is the study by Campelo et al. (*48*), which shows that the formation of a cylindrical tubule is ascribed to the presentation of a polymer concentration gradient. In this work, we show that the elongation of a hemispherical cap membrane into a cylindrical tube may be energetically favorable when critical values of glycocalyx grafting density and glycopolymer length are reached. Additionally, our developed model allows us to investigate the effects of glycocalyx properties (grafting density and length of glycocalyx) on the formation of tubular membrane structures.

Based on our theoretical results, we make the following experimentally relevant predictions. The formation of a cylindrical tubule induced by the glycocalyx depends on its properties and requires to reach the threshold grafting density and glycopolymer length. Two distinct stable states are found, depending on the characteristics of the glycocalyx: a tubular state for high grafting density and long polymer length, and a non-tubular state for low grafting density and short polymer length. Experimentally, this prediction can be tested by conducting experiments to observe the structures induced on Muc1-42TR-expressing cells (*39*) by varying the mucin density. The impact of polymer length on tube formation can also be verified, as it is experimentally feasible to tune the number of tandem repeats (TR) of Muc1 (*39, 76, 77*). For a given glycocalyx grafting density and thickness, the assistance of line tension, spontaneous curvature, and actin force is conducive to tube formation. This prediction could imply that actin polymerization (*16*–*18*) and the action of motor proteins (*19, 20*) on the membrane will make the plasma membrane more favorable to tubulation in living cells. The predictions of our model have implications for understanding the formation of tubular structures generated by the glycocalyx in cells. For example, many unique membrane features in tumor cells are associated with the expression of mucins and hyaluronan on their surface (*35, 78, 79*) and the high incidence of tubular protrusions at the micro- and nanoscale from the cell membrane in various cancer phenotypes (*80*). We expect that the findings from our work will contribute to a better understanding of the role of individual glycocalyx properties in regulating membrane shapes and provide deeper insights into how membrane tubes form under various biophysical conditions.

We note that our present work has some limitations and simplifications, such as prescribing the shape of the membrane as a simplified geometrical scheme, treating the glycocalyx as an uncharged polymer brush network, and neglecting the spatial heterogeneity of the distribution of the glycocalyx polymers on the membrane. Beyond filament polymerization and depolymerization, the attachment of the membrane to the actin cortex may also suppress glycocalyx-mediated membrane morphologies within the cell (*81*–*83*). Future modeling and experimental studies could consider incorporating the diffusion of glycocalyx polymers, changes in the bulk physical properties of the glycocalyx, as well as the effects of membrane-actin cortex attachment and charged polymers to better capture the biological complexity of cell membranes.

## Acknowledgments

This work was supported by NIH R01GM132106, NSF MCB 2327243, and Office of Naval Research N00014-20-1-2469 to P.R. We would like to thank Dr. Emmet Francis for proof reading the manuscript and giving valuable feedback. The authors have no conflicts of interest to declare.

## Supplemental Material

### 1 Derivation of the energy contribution associated with the glycocalyx

The crowding of large glycosylated proteins appears to regulate the shape of the underlying bilayer plasma membrane (*39*). According to polymer physics, the glycocalyx polymers on cell membrane surfaces exhibit two regimes depending on their grafting density, which are the mushroom-like regime and the brush-like regime. In the case of high-density glycocalyx polymers, Shurer et al. (*39*) have reported that the mucins are in the brush-like structure which is able to regulate membrane morphology. To model the influence of the glycocalyx on cell membrane morphology, in our theoretical model, we focus on densely grafted regions of the membrane and therefore model the glycocalyx in the brush regime.

Here, we consider a layer of glycocalyx (brush-like structure) grafted on a membrane with a cylindrical tube and a hemispherical cap geometry, where the tube and cap radii are *R*_*t*_, the tube length is *L*_*t*_, and the thickness of the brush-like structure is *L*_brush_, as shown in Fig. S1. From the viewpoint of coarse-graining, the polymer brush is envisioned as an array of blobs. The size of each blob, *ξ*, at a given position *r*, equals the square root of the local area per chain *s*(*r*), where *r* = *x* + *R*_*t*_ is the radial distance which is defined from the center of the spherical surface or the cylindrical tube surface, and in which *x* is the distance from the membrane surface. Thus, the blob size 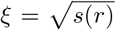 grows as a function of *r*, and the grafting density of glycocalyx polymer brush on the membrane surface can be obtained as *ρ* = 1*/ξ*^2^. Assuming that the layer of glycopolymers is extended non-uniformly but equally in the height, *L*_brush_, then the area per chain at distance *x* from the membrane surface is given by (*52*)

**Figure S1:**
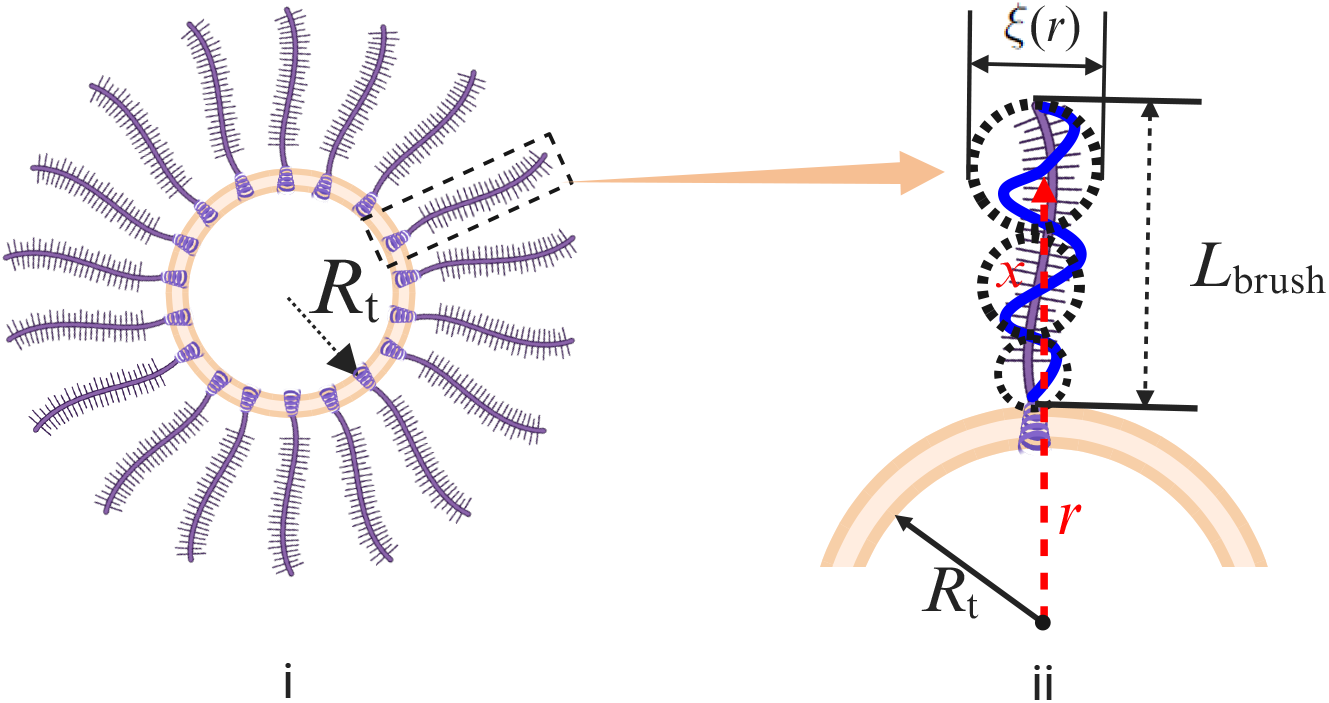
(i) Schematic of the cross-section of a cylindrical tube or a sphere membrane grafted with glycocalyx. (ii) An enlarged schematic illustration of fragments of a polymer chain anchored on a cylinder or sphere with the radius of curvature *R*_*t*_. At a given position *r*, the coarse grained blob (dashed black circle) size in a polymer brush is *ξ*(*r*), and the thickness of the brush is *L*_brush_.

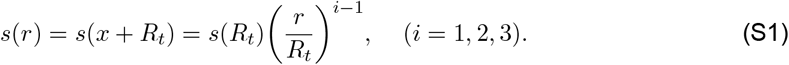

The index *i* = 1, 2, 3 indicates planar, cylindrical, and spherical shaped membranes, respectively. When the membrane is bent, the changes of polymer configuration gives rise to the local extension of the polymer chain. Here the local chain extension at a height *r* is characterized by d*r/*d*n*, where the variable *n* denotes the current monomer. This local extension is related to local density profile of monomers *c*_*p*_(*r*) as (*52*)

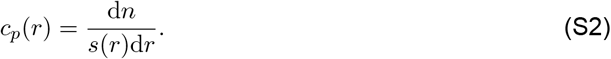

Then, the thickness of the brush, *L*_brush_, is found from the conservation condition (the constraint of conservation of the total number of monomers *N*) (*52*),

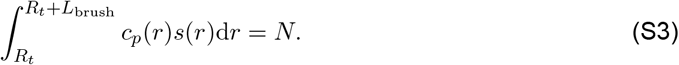

As a result, in the brush regime, since the electrostatic effects are excluded, the energy contribution original from the glycocalyx polymers including two terms (*39, 52*): the elastic energy of the polymer chain 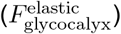 and the free energy caused by the excluded volume interactions of polymer monomers 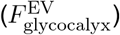 Based on the hypotheses in the main text, according to Ref. (*52*), the elastic energy per chain in the brush can be presented as

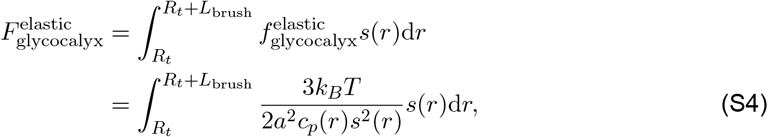

where *a* is the monomer length and 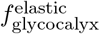 is the elastic energy density of the polymer chain. Within the mean-field approximation, the energy density of the excluded volume interactions (van der Waals interactions) between monomers can be modeled in terms of the virial expansion

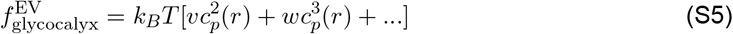

where *υ* and *w* are the second and third virial coefficient, respectively. As a result, the excluded volume interactions between monomers per chain in the brush is given by

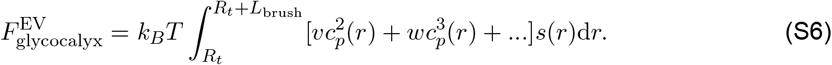

In subsequent analysis, the cubic and higher terms are neglected. Therefore, the sum of Eq. (S4) and Eq. (S6) yields the energy contribution of glycocalyx polymers

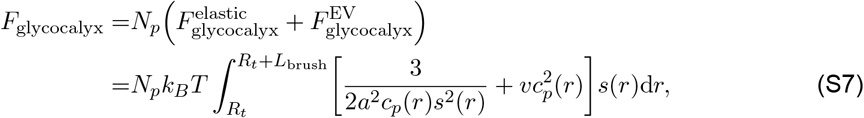

where *N*_*p*_ is the number of polymer chains grafted on the membrane.

In order to calculate the free energy, we need to further determine the local concentration of monomers *c*_*p*_(*r*) and the brush thickness *L*_brush_. On a planar membrane surface, the planar brush area per chain, *s*(*r*), is constant, i.e., *s*(*r*) = *s*. Minimizing the free energy of a planar brush F_glycocalyx_ with respect to *c*_*p*_ by using 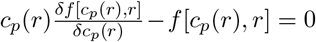 leads to the equilibrium polymer concentration 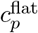 (see Ref. (52) for detailed steps)

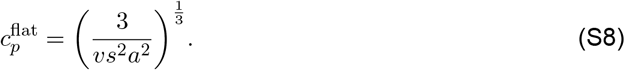

Here, 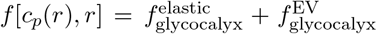 Using the conservation condition defined by Eq. (S3) yields the brush thickness 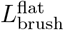 (see Ref. (52) for detailed steps)

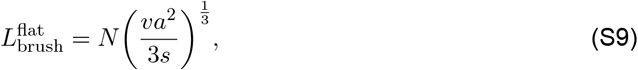

where we can see that the thickness of polymer brush is proportional to the total number of monomers *N*. Hereafter, in our model, we use the number of monomers, *N*, to capture the length of the polymer. Note that the relationship for 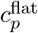 and 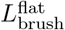 are special cases of electrically neutral brushes as described in (*52*). Substituting Eq. (S8) and Eq. (S9) into Eq. (S7) yields the energy contribution of glycocalyx polymers on a planar membrane surface

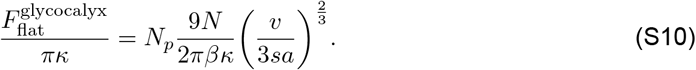

Thus, even for a flat membrane, the energy contribution by the glycocalyx is directly proportional to the extent of grafting *N*_*p*_ and the length of the polymer brush *N*.

On a spherical or a cylindrical membrane surface, the corresponding monomer density profile *c*_*p*_(*r*) and brush thickness *L*_brush_ are, respectively, expressed as (see Ref. (*52*) for a detailed calculation)

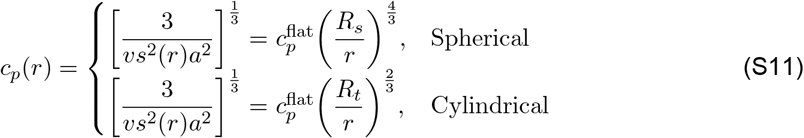

and

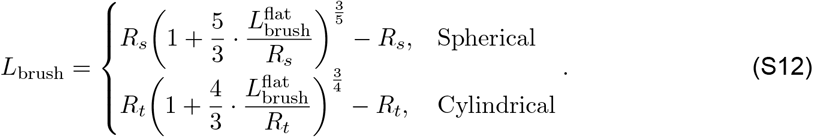

Similarly, substituting Eq. (S11) and Eq. (S12) into Eq. (S7) leads to the energy contribution of glycocalyx polymers on a spherical or a cylindrical membrane surface as

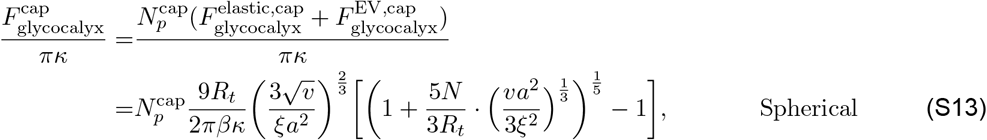

and

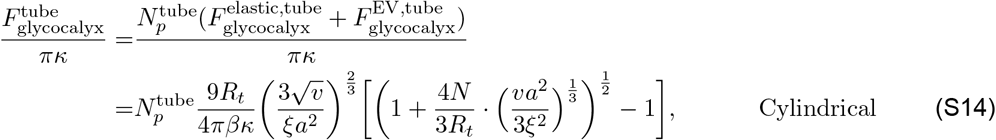

where 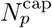 and 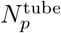 are the number of polymer chains that grafted on the spherical cap and the cylindrical tube. Combining Eqs. (S13) and (S14), and using the relation *χ* = *L*_*t*_*/R*_*t*_ defined in the main text reduces to Eq. (7) in the main text.

### 2 Contributions from the different energy components

To reveal the mechanisms of tube formation regulated by glycocalyx, different energy components contribute to the total free energy profiles in Fig. 2 are plotted as a function of shape parameter *χ*, as shown in Fig. S2. Figure. S2 demonstrates that membrane bending is governed by a balance among these energy players.

**Figure S2:**
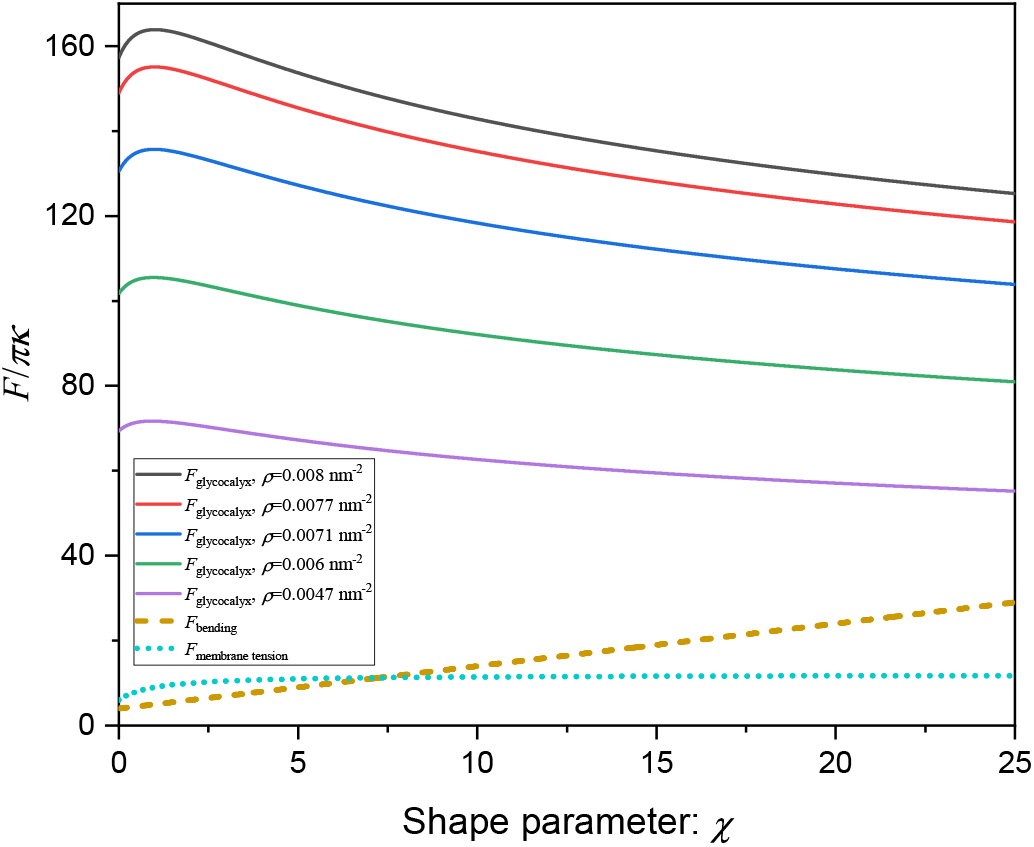
Different types of energy profiles including energy contribution associated with glycocalyx *F*_glycocalyx_, bending energy *F*_bending_, and tension energy *F*_membrane tension_ as a function of the shape parameter *χ* for different grafting densities.

### 3 Phase diagrams on the (*κ*–*σ*), (*λ*–*c*_0_), and (*f* –*c*_0_) planes

To gain more insight into the effects of the membrane properties and spontaneous curvature, line tension, and actin force on tube formation, three phase diagrams on the *κ*–*σ, λ*–*c*_0_, and *f* –*c*_0_ plane are constructed, as shown in Fig. S3. Figure S3(a) shows that forming a tube is more favorable under lower membrane bending rigidity and membrane tension. Figure S3(b) and (c) confirmed that longer and thinner tube can be formed with the assistance of line tension, spontaneous curvature, and actin force.

**Figure S3:**
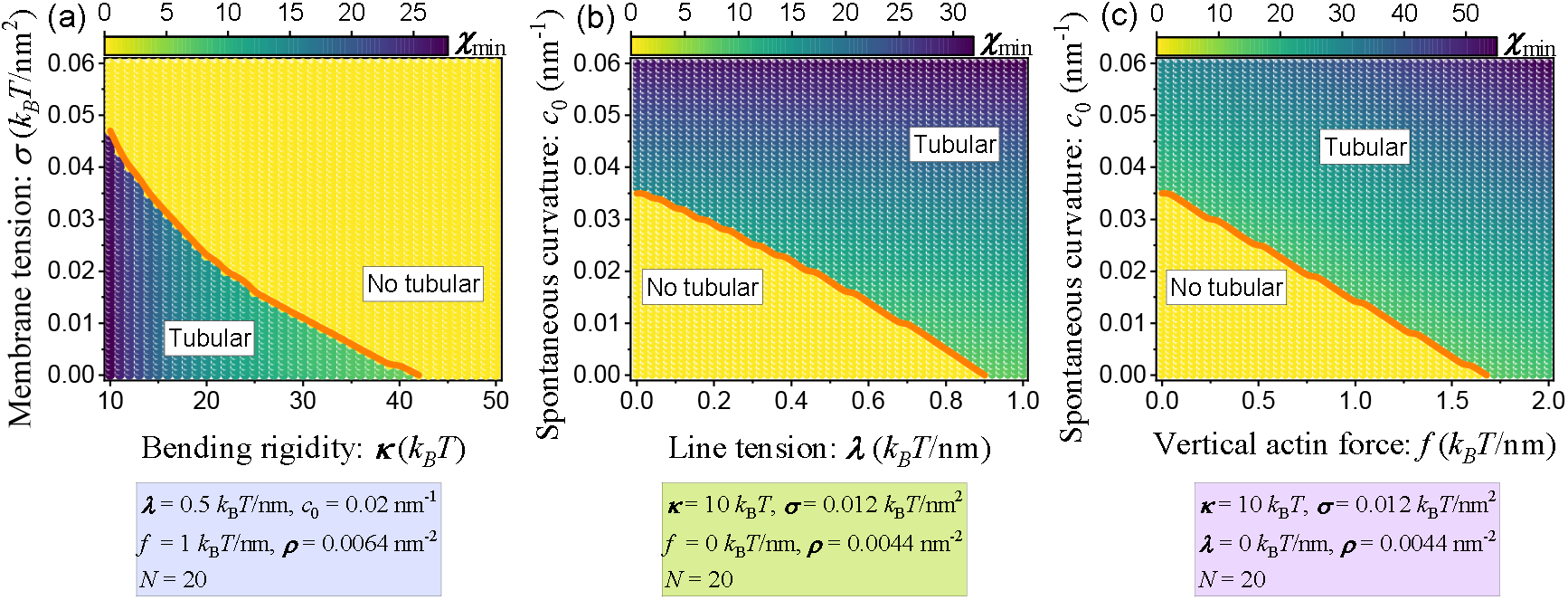
(a) Heatmap of optimal shape parameter *χ*_min_ as a function of membrane bending rigidity *κ* and membrane tension *σ*. (b) Contour plot of optimal shape parameter *χ*_min_ as a function of line tension *λ* and spontaneous curvature *c*_0_, where the color bar represents the magnitude of the optimal shape parameter. (c) A two-dimensional phase diagram on the (*f* –*c*_0_) plane characterizes the interrelated effects of vertical actin force and spontaneous curvature on the membrane shape.

## Notes

### Competing Interest Statement

The authors have declared no competing interest.

## References

1. C. Lee, L. B. Chen, Cell 54, 37–46, DOI 10.1016/0092-8674(88)90177-8 (1988).

2. L. Westrate, J. Lee, W. Prinz, G. Voeltz, Annu. Rev. Biochem. 84, 791–811, DOI 10.1146/annurev-biochem-072711-163501 (2015).

3. H. Mollenhauer, D. Morré, Histochem. Cell Biol. 109, 533–543, DOI 10.1007/s004180050253 (1998).

4. M. A. De Matteis, A. Luini, Nat. Rev. Mol. Cell Biol. 9, 273–284, DOI 10.1038/nrm2378 (2008).

5. T. G. Frey, C. A. Mannella, Trends Biochem. Sci. 25, 319–324, DOI 10.1016/S0968-0004(00)01609-1 (2000).

6. C. A. Mannella, Biochim. Biophys. Acta 1763, 542–548, DOI 10.1016/j.bbamcr.2006.04.006 (2006).

7. C. Wang et al., Proc. Natl. Acad. Sci. U. S. A. 116, 15817–15822, DOI 10.1073/pnas.1905924116 (2019).

8. P. K. Mattila, P. Lappalainen, Nat. Rev. Mol. Cell Biol. 9, 446–454, DOI 10.1038/nrm2406 (2008).

9. T. Hong et al., Nat. Med. 20, 624–632, DOI 10.1038/nm.3543 (2014).

10. A. Mahapatra, C. Uysalel, P. Rangamani, J. Membr. Biol. 254, 273–291, DOI 10.1007/s00232-020-00164-9 (2021).

11. Z. Feng, C.-h. Yu, Proc. Natl. Acad. Sci. U. S. A. 118, e2017645118, DOI 10.1073/pnas.2017645118 (2021).

12. R. Waugh, Biophys. J. 38, 29–37, DOI 10.1016/S0006-3495(82)84527-X (1982).

13. E. Evans, H. Bowman, A. Leung, D. Needham, D. Tirrell, Science 273, 933–935, DOI 10.1126/science.273.5277.933 (1996).

14. D. Raucher, M. P. Sheetz, Biophys. J. 77, 1992–2002, DOI 10.1016/S0006-3495(99)77040-2 (1999).

15. V. Heinrich, R. Waugh, Ann. Biomed. Eng. 24, 595–605, DOI 10.1007/BF02684228 (1996).

16. H. Miyata, H. Hotani, Proc. Natl. Acad. Sci. U. S. A. 89, 11547–11551, DOI 10.1073/pnas.89.23.11547 (1992).

17. H. Miyata, S. Nishiyama, K.-i. Akashi, K. Kinosita, Proc. Natl. Acad. Sci. U. S. A. 96, 2048–2053, DOI 10.1073/pnas.96.5.2048 (1999).

18. J. Weichsel, P. L. Geissler, PLoS Comput. Biol. 12, 1–13, DOI 10.1371/journal.pcbi.1004982 (2016).

19. O. Campàs et al., Biophys. J. 94, 5009–5017, DOI 10.1529/biophysj.107.118554 (2008).

20. W. Du et al., Dev. Cell 37, 326–336, DOI 10.1016/j.devcel.2016.04.014 (2016).

21. I. Derényi, F. Jülicher, J. Prost, Phys. Rev. Lett. 88, 238101, DOI 10.1103/PhysRevLett.88.238101 (2002).

22. A. Callan-Jones, P. Bassereau, Curr. Opin. Solid State Mater. Sci. 17, 143–150, DOI 10.1016/j.cossms.2013.08.004 (2013).

23. A. Frost, V. M. Unger, P. De Camilli, Cell 137, 191–196, DOI 10.1016/j.cell.2009.04.010 (2009).

24. A. Mahapatra, P. Rangamani, Soft Matter 19, 4345–4359, DOI 10.1039/D2SM01676A (2023).

25. N. Walani, J. Torres, A. Agrawal, Phys. Rev. E 89, 062715, DOI 10.1103/PhysRevE.89.062715 (2014).

26. K. Xiao, C.-X. Wu, R. Ma, Phys. Rev. Res. 5, 023176, DOI 10.1103/PhysRevResearch.5.023176 (2023).

27. C. L. Hattrup, S. J. Gendler, Annu. Rev. Physiol. 70, 431–457, DOI 10.1146/annurev.physiol.70.113006.100659 (2008).

28. Y. Jung et al., Proc. Natl. Acad. Sci. U. S. A. 113, E5916–E5924, DOI 10.1073/pnas.1605399113 (2016).

29. G. Kesavan et al., Cell 139, 791–801, DOI 10.1016/j.cell.2009.08.049 (2009).

30. M. Kesimer et al., Mucosal Immunol. 6, 379–392, DOI 10.1038/mi.2012.81 (2013).

31. S. Makabe, T. Naguro, T. Stallone, Microsc. Res. Tech. 69, 436–449, DOI 10.1002/jemt.20303 (2006).

32. S. P. Evanko, M. I. Tammi, R. H. Tammi, T. N. Wight, Adv. Drug Delivery Rev. 59, 1351–1365, DOI 10.1016/j.addr.2007.08.008 (2007).

33. B. Button et al., Science 337, 937–941, DOI 10.1126/science.1223012 (2012).

34. J. Richard Bennett et al., J. Histochem. Cytochem. 49, 67–77, DOI 10.1177/002215540104900107 (2001).

35. V. Koistinen et al., Exp. Cell Res. 337, 179–191, DOI 10.1016/j.yexcr.2015.06.016 (2015).

36. J. C.-H. Kuo, J. G. Gandhi, R. N. Zia, M. J. Paszek, Nat. Phys. 14, 658–669, DOI 10.1038/s41567-018-0186-9 (2018).

37. S. Weinbaum, J. M. Tarbell, E. R. Damiano, Annu. Rev. Biomed. Eng. 9, 121–167, DOI 10.1146/annurev.bioeng.9.060906.151959 (2007).

38. L. Möckl, Front. Cell Dev. Biol. 8, DOI 10.3389/fcell.2020.00253 (2020).

39. C. R. Shurer et al., Cell 177, 1757–1770.e21, DOI 10.1016/j.cell.2019.04.017 (2019).

40. W. Helfrich, Z. Naturforsch. C 28, 693–703, DOI 10.1515/znc-1973-11-1209 (1973).

41. E. Evans, Biophys. J. 14, 923–931, DOI 10.1016/S0006-3495(74)85959-X (1974).

42. C. Hiergeist, R. Lipowsky, J. Phys. II France 6, 1465–1481, DOI 10.1051/jp2:1996142 (1996).

43. R. Lipowsky, Europhys. Lett. 30, 197, DOI 10.1209/0295-5075/30/4/002 (1995).

44. M. Breidenich, R. R. Netz, R. Lipowsky, Europhys. Lett. 49, 431, DOI 10.1209/epl/i2000-00167-2 (2000).

45. T. Bickel, C. Jeppesen, C. Marques, Eur. Phys. J. E 4, 33–43, DOI 10.1007/s101890170140 (2001).

46. Y. W. Kim, W. Sung, Phys. Rev. E 63, 041910, DOI 10.1103/PhysRevE.63.041910 (2001).

47. F. Campelo, A. Hernández-Machado, Phys. Rev. Lett. 99, 088101, DOI 10.1103/PhysRevLett.99.088101 (2007).

48. F. Campelo, A. Hernández–Machado, Phys. Rev. Lett. 100, 158103, DOI 10.1103/PhysRevLett.100.158103 (2008).

49. M. Werner, J. U. Sommer, Eur. Phys. J. E 31, 383–392, DOI 10.1140/epje/i2010-10576-4 (2010).

50. S. Kutti Kandy, R. Radhakrishnan, Biophys. J. 121, 3674–3683, DOI 10.1016/j.bpj.2022.05.031 (2022).

51. K. Xiao, S. Park, J. C. Stachowiak, P. Rangamani, biorxiv, DOI 10.1101/2024.09.07.611813 (2024).

52. E. B. Zhulina, T. M. Birshtein, O. V. Borisov, Eur. Phys. J. E 20, 243–256, DOI 10.1140/epje/i2006-10013-5 (2006).

53. A. Mogilner, G. Oster, Biophys. J. 71, 3030–3045, DOI 10.1016/S0006-3495(96)79496-1 (1996).

54. A. Mogilner, G. Oster, Biophys. J. 84, 1591–1605, DOI 10.1016/S0006-3495(03)74969-8 (2003).

55. I. Raote et al., eLife 9, e59426, DOI 10.7554/eLife.59426 (2020).

56. N. Walani, J. Torres, A. Agrawal, Proc. Natl. Acad. Sci. U. S. A. 112, E1423–E1432, DOI 10.1073/pnas.1418491112 (2015).

57. R. Ma, J. Berro, Biophys. J. 120, 1625–1640, DOI 10.1016/j.bpj.2021.02.033 (2021).

58. R. Lipowsky, Biophys. J. 64, 1133–1138, DOI 10.1016/S0006-3495(93)81479-6 (1993).

59. J. Liu, M. Kaksonen, D. G. Drubin, G. Oster, Proc. Natl. Acad. Sci. U. S. A. 103, 10277–10282, DOI 10.1073/pnas.0601045103 (2006).

60. J.-M. Allain, C. Storm, A. Roux, M. B. Amar, J.-F. Joanny, Phys. Rev. Lett. 93, 158104, DOI 10.1103/PhysRevLett.93.158104 (2004).

61. L. Foret, Eur. Phys. J. E 37, DOI 10.1140/epje/i2014-14042-1 (2014).

62. Z. Shi, T. Baumgart, Nat. Commun. 6, DOI 10.1038/ncomms6974 (2015).

63. R. Phillips, T. Ursell, P. Wiggins, P. Sens, Nature 459, 379–385, DOI 10.1038/nature08147 (2009).

64. T. Baumgart, S. Hess, W. Webb, Nature 425, 821–824, DOI 10.1038/nature02013 (2003).

65. R. Lipowsky, J. Phys. II France 2, 1825–1840, DOI 10.1051/jp2:1992238 (1992).

66. D. Bracha, E. Karzbrun, G. Shemer, P. A. Pincus, R. H. Bar-Ziv, Proc. Natl. Acad. Sci. U. S. A. 110, 4534–4538, DOI 10.1073/pnas.1220076110 (2013).

67. L. Foret, P. Sens, Proc. Natl. Acad. Sci. U. S. A. 105, 14763–14768, DOI 10.1073/pnas.0801173105 (2008).

68. J. Paturej, S. S. Sheiko, S. Panyukov, M. Rubinstein, Sci. Adv. 2, e1601478, DOI 10.1126/sciadv.1601478 (2016).

69. J. G. Gandhi, D. L. Koch, M. J. Paszek, Biophys. J. 116, 694–708, DOI 10.1016/j.bpj.2018.12.023 (2019).

70. D. R. Kovar, T. D. Pollard, Proc. Natl. Acad. Sci. U. S. A. 101, 14725–14730, DOI 10.1073/pnas.0405902101 (2004).

71. C. Peskin, G. Odell, G. Oster, Biophys. J. 65, 316–324, DOI 10.1016/S0006-3495(93)81035-X (1993).

72. T. D. Pollard, G. G. Borisy, Cell 112, 453–465, DOI 10.1016/S0092-8674(03)00120-X (2003).

73. S. M. Block, C. L. Asbury, J. W. Shaevitz, M. J. Lang, Proc. Natl. Acad. Sci. U. S. A. 100, 2351–2356, DOI 10.1073/pnas.0436709100 (2003).

74. M. J. Footer, J. W. J. Kerssemakers, J. A. Theriot, M. Dogterom, Proc. Natl. Acad. Sci. U. S. A. 104, 2181–2186, DOI 10.1073/pnas.0607052104 (2007).

75. K. Xiao, R. Ma, C.-X. Wu, Phys. Rev. E 106, 044411, DOI 10.1103/PhysRevE.106.044411 (2022).

76. C.-H. Lu et al., Nat. Commun. 13, 3093, DOI 10.1038/s41467-022-30610-2 (2022).

77. S. Gollapudi et al., Proc. Natl. Acad. Sci. U.S.A. 120, e2215815120, DOI 10.1073/pnas.2215815120 (2023).

78. D. W. Kufe, Nat. Rev. Cancer 9, 874–885, DOI 10.1038/nrc2761 (2009).

79. E. A. Turley, D. K. Wood, J. B. McCarthy, Cancer Res. 76, 2507–2512, DOI 10.1158/0008-5472.CAN-15-3114 (2016).

80. R. Xu et al., Nat. Rev. Clin. Oncol. 15, 617–638, DOI 10.1038/s41571-018-0036-9 (2018).

81. E. S. Welf et al., Dev. Cell 55, 723–736.e8, DOI 10.1016/j.devcel.2020.11.024 (2020).

82. A. Diz-Muñoz et al., PLoS Biol. 8, 1–12, DOI 10.1371/journal.pbio.1000544 (2010).

83. A. Paraschiv et al., Biophys. J. 120, 598–606, DOI 10.1016/j.bpj.2020.12.028 (2021).

